# Redefining Parkinson’s Disease by Dysregulated Genetic Networks in Distinct Cell Types

**DOI:** 10.1101/2025.09.09.674964

**Authors:** Jane Yang, Arianna Colini Baldeschi, JungHo Kong, Nicole Mattson, Erin Schiksnis, Sarah Wright, SungJoon Park, InGoo Lee, Laurent Roybon, Trey Ideker

## Abstract

Parkinson’s disease (PD) is classically linked to dopaminergic neuron loss, but emerging evidence suggests broader cellular involvement. Here we show that PD risk variants converge on distinct molecular networks across specific brain cell types, enabling stratification of patients into six subgroups: dopaminergic, oligodendrocyte progenitor cells (O), excitatory (E), dopaminergic/excitatory, dopaminergic/oligodendrocyte and other. While all subgroups exhibit motor symptoms, the E-group individuals also display more severe non-motor symptoms, including dementia, hyposmia, and REM sleep behavior disorder. The O-group individuals exhibit reduced myelin integrity, as demonstrated by diffusion tensor imaging, implicating *NRG6* (formerly *C1orf56*), a previously uncharacterized high-risk PD gene. We show that *NRG6* encodes a conserved epidermal growth factor-like domain structurally and functionally analogous to neuregulin-1, which is critical for oligodendrocyte development and myelination. These findings redefine the cellular architecture of PD vulnerability and identify neuregulin-like signaling in oligodendrocytes as a potential contributor to non-motor symptoms.

## INTRODUCTION

Parkinson’s disease (PD) is a progressive neurodegenerative disorder which principally affects movement, with approximately 10 million people affected worldwide^1^. Primary motor symptoms result from the degeneration of dopaminergic (DA) neurons in the substantia nigra, which are particularly vulnerable because of their high pacemaking activity^2^ and dense axonal arborization^3^ which require elevated respiration leading to oxidative stress^4^. Other cell types have also been implicated^5,6^, however, with white matter (non-neuronal) impairment shown to precede DA degeneration^7^. Understanding the role of these other cell types could lead to new treatments beyond current DA-targeting drugs, which tend to address motor symptoms but not disease progression and bring adverse side effects, such as nausea, drowsiness, and neuropsychiatric manifestations^8^.

The development of single-cell and single-nuclei resolution mRNA sequencing has enabled profiling of transcriptomic changes in specific cell types from post-mortem human brain tissues^9,10^. Analysis of these data has shown that genes highly expressed in dopaminergic neurons are significantly enriched for common risk variants^11^. Beyond dopaminergic neurons, Smajić et al.^12^ observed increased microglial and astrocyte activation, with fewer oligodendrocytes, and Martirosyan et al.^13^ reported an overall increase in glial and T cells. While both studies highlight a role for non-neuronal cells in PD via immunomodulation, Agarwal et al.^11^ found no association between PD risk and genes expressed in astrocytes or microglia. These conflicting results raise the question of whether the observed changes in non-neuronal cells are a cause of, or a response to, degeneration.

Here we study the principal molecular networks, and their associated cell types, that underlie genetic risk for PD (**Fig. 1a**). This analysis suggests a redefinition of this disease as a collection of six subtypes, each involving distinct networks, which also stratify key disease symptoms (**Fig. 1b**). Among these, genetic circuits in excitatory neurons and oligodendrocyte precursor cells (OPCs) emerge as key drivers of PD risk. Notably, we implicate *NRG6* (formerly *C1orf56*), a functionally uncharacterized gene, in oligodendrocyte differentiation, pointing to a glial mechanism of disease independent of dopaminergic loss (**Fig. 1c**).

**Figure 1.**
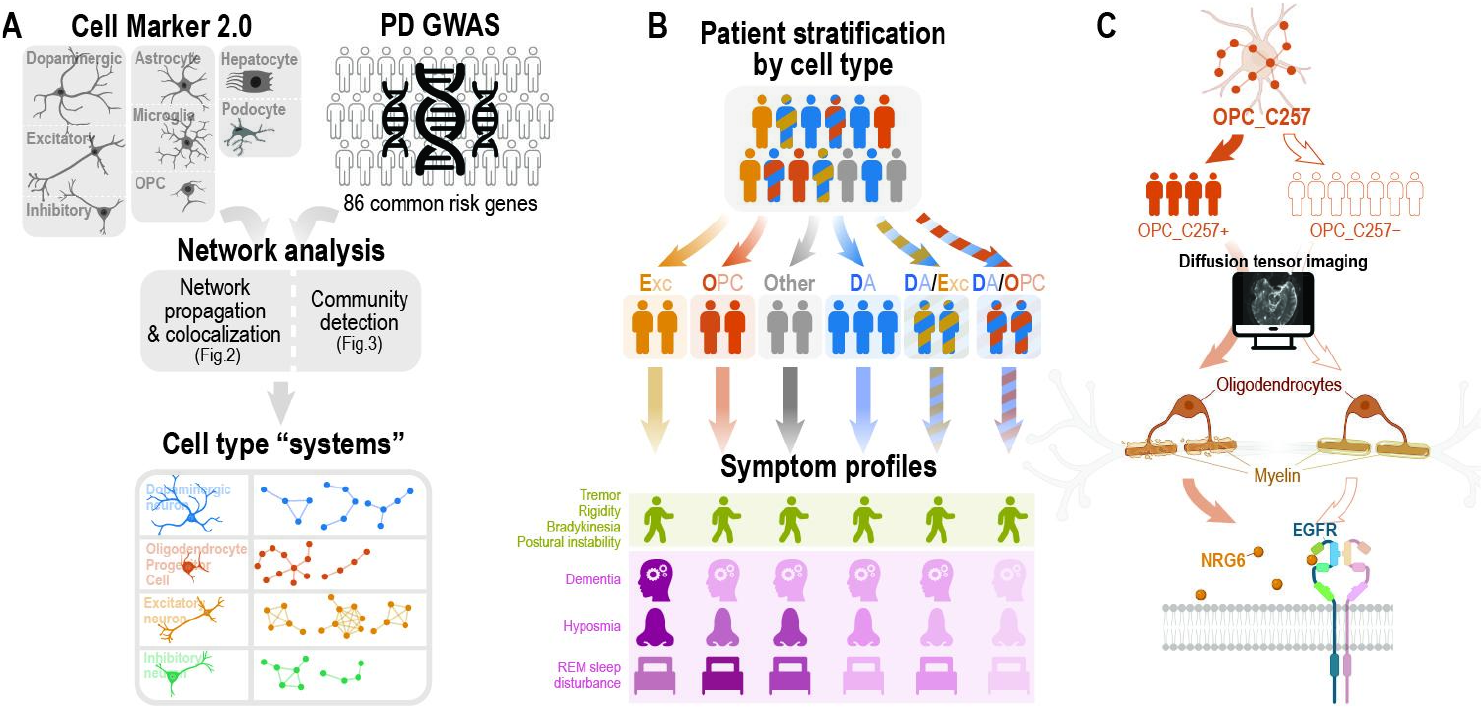
Classifying PD patients by genetic networks in distinct cell types. (**a**) Common PD risk genes are integrated with cell-type molecular markers to identify dysregulated molecular networks, or “systems,” affected in each cell type. (**b**) Using the systems framework established in A, PD cases from an independent cohort are classified as E (excitatory), O (oligodendrocyte progenitor cell; OPC), D (dopaminergic), D/E (dopaminergic/excitatory), D/O (dopaminergic/OPC), or other. The six groups are similar in some aspects but differ in non-motor symptoms. (**c**) The previously unidentified “OPC_C257” system is confirmed to affect myelin integrity through diffusion tensor imaging and, for the first time, implicates *NRG6* (formerly *C1orf56*) in oligodendrocyte biology through structural and functional parallels with neuregulin-1.

## RESULTS

### Convergence of PD risk genes and brain cell type markers within genetic networks

We obtained 90 genome-wide significant variants^14^ and mapped each to the nearest gene to define a non-overlapping set of 86 common PD risk genes. In parallel, we obtained gene markers specific to six brain cell types, including 18 DA neuron, 28 excitatory neuron, 28 inhibitory neuron, 96 OPC, 328 microglia, and 394 astrocyte markers^15^ (**Fig. 2a**). We also obtained 32 hepatocyte and 15 podocyte markers, representing two cell types outside the brain to be used as negative controls. To study these genes in a network context, we accessed HumanNet-XC^16^, a comprehensive gene interaction database which integrates various types of evidence including protein-protein interactions, genetic interactions, gene co-expression and literature-based associations– yielding a network of 1,125,494 functional relationships among 18,593 human genes.

**Figure 2.**
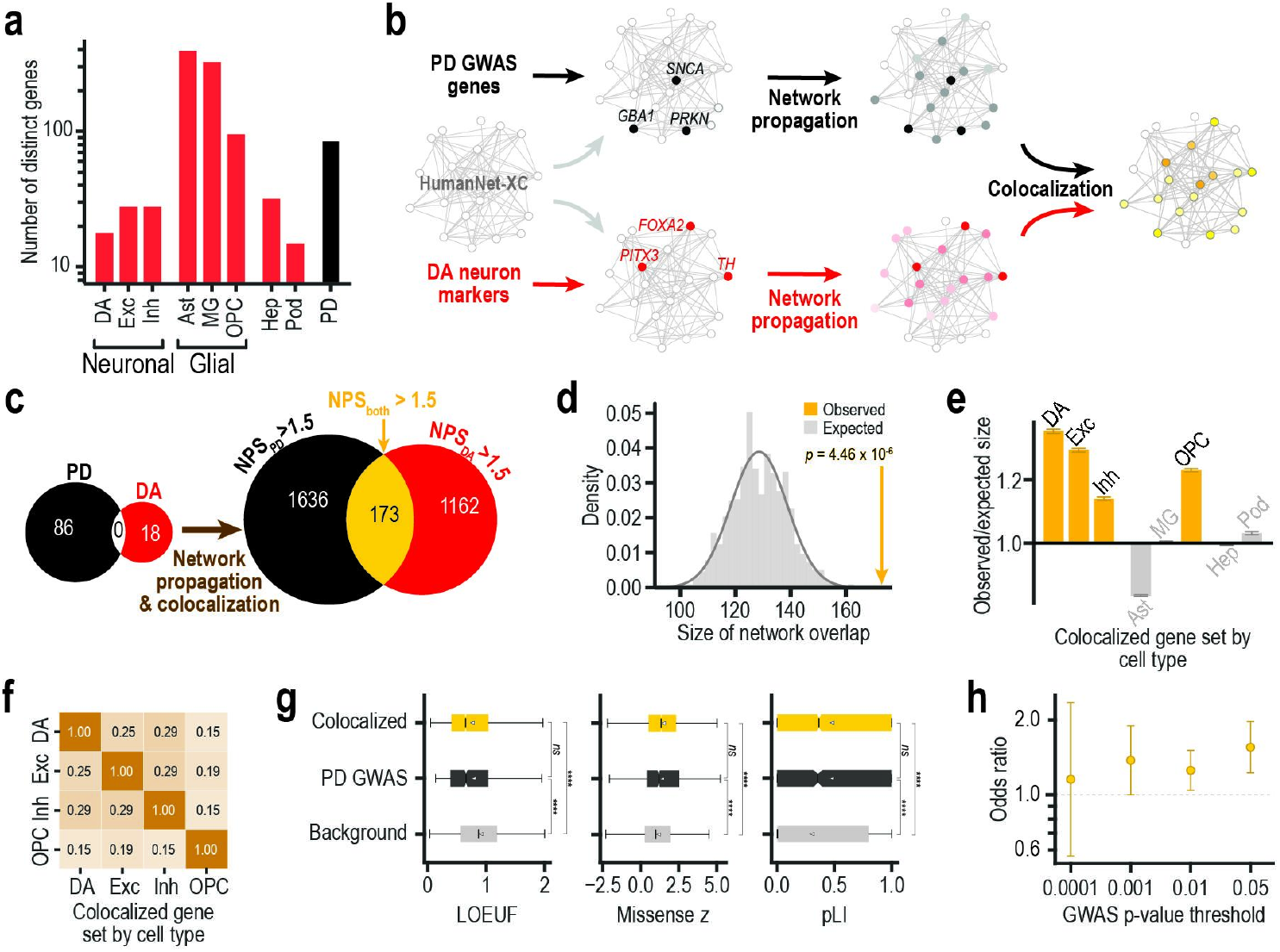
Convergence of cell markers with common risk genes on molecular networks. (**a**) Number of distinct gene markers for brain cell types (red) and number of Parkinson’s Disease common risk genes (black). (**b**) Toy illustration of network propagation and colocalization using DAn markers as an example. Sets of common risk genes and gene markers (left) are propagated separately in a molecular interaction network (center). Colocalized genes are near both common risk genes and cell markers in the network (right). Subsets of the network and gene sets are shown for illustrative purposes. (**c**) Venn diagrams showing gene overlap between common risk genes and DA neuron markers before (left) and after network propagation and colocalization (right). The colocalized gene set (yellow) is defined based on network proximity scores (NPS) as the set of genes with all of NPS_PD_ >1.5, NPS_DA_ > 1.5 and NPS_both_ >1.5. (**d**) Observed (yellow arrow) vs. expected size (gray distribution, 10,000 permutations) of the colocalized gene set for DAn, *p* = 4.46 × 10^-6^ (*Z* test). (**e**) Observed-to-expected ratio of colocalized gene sets for all eight cell types (yellow, *p* < 0.05 by *Z* test). Error bars indicate 95% confidence intervals. (**f**) Similarity of colocalized gene sets across cell-type pairs (Jaccard similarity index). (**g**) Distributions of LOEUF, missense Z, and pLI for background, PD GWAS, and colocalized genes. Boxes indicate the interquartile range (IQR); the notch marks the median; the white marker denotes the mean; the whiskers extend to 1.5×IQR. LOEUF, loss-of-function observed/expected upper-bound fraction; missense Z, standardized residual for missense variation; pLI; probability of LoF intolerance. (**h**) Enrichment of the colocalized genes among genes implicated by the Nalls et al. PD GWAS (summary statistics), defined as nearest genes to SNPs meeting each GWAS p-value threshold. Vertical error bars indicate 95% confidence intervals; the grey dashed line marks OR = 1. **** *p* < 0.0001; *ns* = non-significant (two-tailed Mann-Whitney *U* test). PD, Parkinson’s Disease; DAn, dopaminergic neuron; Exc, excitatory neuron; Inh, inhibitory neuron; OPC, oligodendrocyte progenitor cell.

Next, we provided the PD risk genes as ‘seeds’ for network propagation^17^, a probabilistic approach that spreads signal from each seed outward through the network, transmitting high proximity scores to nearby genes (also known as heat diffusion, **Methods**). Network propagation was also performed using each set of cell-type markers as seeds. We then integrated these network propagation results, yielding sets of ‘co-localized’ genes having close network proximity to both PD risk genes and markers of a particular cell type.

To test whether our approach successfully recapitulates PD-specific biology, we first examined DA neurons (DAn) as a positive control (**Fig. 2b**), given their selective vulnerability in PD. The network proximity of each human gene to the set of PD risk genes, and separately to the set of DAn markers, was calculated using network propagation. These proximity scores were then thresholded to extract a PD-DA colocalized set of genes (**Methods, Fig. 2c**), which was significantly larger than expected by chance (**Fig. 2d**). In addition to DAn, we found that cell-type specific markers of excitatory neurons, inhibitory neurons, and OPCs demonstrated significant network colocalization with PD risk genes, while astrocytes, microglia, and the two negative controls did not (**Fig. 2e**). These colocalized gene sets were largely different from cell type to cell type (**Fig. 2f**).

These genes were also more constrained than background genes and as constrained as GWAS genes (**Fig. 2g**), implying that they are functionally important and cannot tolerate changes in gene dosage. They were significantly enriched for nominal GWAS genes, excluding seed genes, with enrichment increasing under more permissive p-value thresholds (**Fig. 2h**; p < 1×10^-4^: OR = 1.2, 95% CI = 0.6–2.4; p < 1×10^-3^: OR = 1.4, 95% CI = 1.0–1.9; p < 1×10^-2^: OR = 1.3, 95% CI = 1.0–1.5; p < 0.05: OR = 1.6, 95% CI = 1.2–2.0). Together, these findings indicate that our method captures the polygenic architecture of PD driven by common variants of modest effects.

### Identification of PD-associated molecular systems in distinct cell types

We found that each set of colocalized genes could be subdivided into a series of densely interacting network communities (**Methods, Fig. 3a** shows the example of DAn). The resulting 89 communities, henceforth called network “systems,” included 21 DAn, 24 excitatory neuron, 19 inhibitory neuron, and 25 OPC systems, respectively. These systems ranged in size from 4 to 17 genes and were largely distinct from one another (**Fig. 3b**).

**Figure 3.**
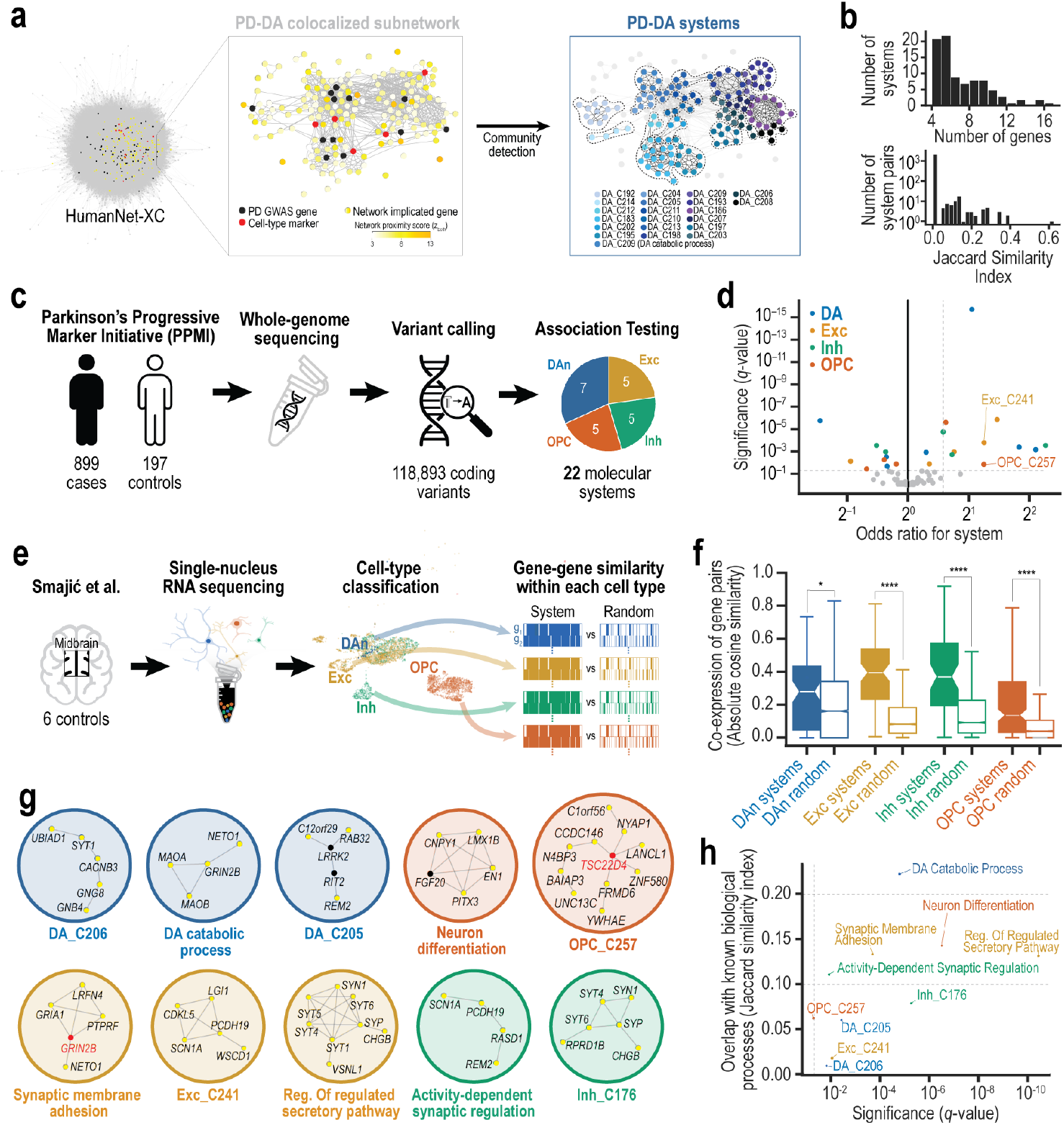
Identification of PD-risk systems regulated within distinct cell types. (**A**) Visualization of 21 DAn systems, or gene communities, detected from the PD-DA colocalized gene set. Left, HumanNet-XC showing PD GWAS genes (black), DAn markers (red), colocalized genes (yellow), and others (gray). Middle, PD-DA colocalized subnetwork extracted based on proximity between PD GWAS genes and DAn markers (z_both_), with scores ranging from low (light yellow; distant) to high (dark yellow; close). Right, community detection applied to the PD-DA subnetwork revealing 21 DA systems (DA_C192 - DA_C214), each represented by a distinct color. (**B**) Distributions of system sizes (top, number of genes per system) and overlaps between systems (bottom, Jaccard similarity indices of all possible system pairs). (**C**) Workflow depicting the replication of systems in the Parkinson’s Progressive Marker Initiative (PPMI) cohort. Whole genome sequencing data for cases and controls are analyzed for coding variants, used to test each system for significant association between variant burden and PD risk. The pie chart shows the cell types of the 22 replicated systems. (**D**) Volcano plot showing the significance versus strength of PD association for each system. The horizontal dashed line indicates an FDR cutoff of *q* = 0.05. Odds ratios > 1 indicate a stronger association with cases, whereas odds ratios < 1 indicate a stronger association with controls. The vertical dashed line indicates the odds ratio cutoff (OR = 1.5) used to select the top ten PD-risk systems. (**E**) Workflow illustrating the assessment of gene functional relevance within molecular systems using single-nucleus RNA sequencing data from midbrain samples by Smajić et al. (**F**) Degree of co-expression of gene pairs (y axis), with genes selected from network systems of each cell type (x axis, solid color boxplots) versus other genes (outline boxplots, 1,000 randomly selected gene pairs per boxplot), computed separately for DAn (n = 32), Exc (n = 1,821), Inh (n = 1,085), and OPCs (n = 1,538). * *p* < 0.05 and **** *p* < 0.0001 (two-tailed Mann-Whitney *U* test). (**G**) Gene networks of the ten PD-risk systems having an odds ratio > 1.5. (**H**) Overlap of PD-risk systems with GO Biological Processes (y axis), selected based on enrichment significance (x axis). The vertical dashed line indicates an FDR threshold of *q* = 0.05. Systems with Jaccard similarity index (JSI) > 0.2 are considered closely aligned, JSI > 0.1 weakly aligned, and otherwise not aligned (JSI thresholds shown as horizontal dashed lines). Systems not aligned (JSI ≤ 0.1) are assigned systematic names. PD, Parkinson’s Disease; DAn, dopaminergic neuron; Exc, excitatory neuron; Inh, inhibitory neuron; OPC, oligodendrocyte progenitor cell.

Prior to further analysis, we sought to validate the network systems against an independent cohort obtained from the Parkinson’s Progressive Marker Initiative (PPMI), a longitudinal study tracking participants over time with whole-genome sequencing as well as repeated clinical, imaging, and biological assessments (**Fig. 3c**). As an orthogonal validation strategy, we used variant burden analysis to test whether these network-defined systems carry significantly higher burden in PD cases than in non-PD controls. This design integrates complementary aspects of the polygenic architecture of PD (**Fig. 2h**): common noncoding variation, which highlights genomic regions associated with PD risk, and rare coding variation, which pinpoints genes within these systems that may directly impact protein function. In brief, we first computed the system-wide burden scores of rare coding variants with potential functional consequences (frameshift, splice-site/region, missense, nonsense, and nonstop) for each of the 89 systems, without additional filtering by predicted deleteriousness (**Methods**). We then used this system-wide burden to model case/control status across individuals, yielding 22 systems with significant PD association (FDR < 0.05, **Fig. 3d, Table S1**). To confirm their biological plausibility, we used single-nucleus RNA sequencing data from an independent cohort of non-PD controls^12^ to evaluate gene expression coherence (**Fig. 3e**), and found consistent co-regulation within the relevant cell types (**Fig. 3f**).

Of the validated systems, we noted 10 in particular (**Fig. 3g**) in which variants were associated with substantial risk for PD (odds ratio ≥ 1.5, **Fig. 3d**), whereas others were protective (10 systems) or of lesser effect (2 systems). Functional analysis indicated that only one of these systems was closely aligned with well-known biological processes, while the remaining either corresponded only weakly to known biology or did not align at all (**Fig. 3h**). The lack of alignment suggests that these systems may represent unique functions that are not well captured by existing pathway annotations.

### Stratification of patients by impacted network systems

Systems conferring PD risk were used to stratify individual cases into six robust subgroups based on variant patterns (**Fig. 4a**; see **Fig. S1** for robustness analysis). These PD subgroups were then assigned alphabetic codes based on the cell types associated with their altered systems. These were E (excitatory neurons and not DA), D/E (DA and excitatory), O (OPC only), D/O (DA and OPC), and D (DA only) subgroups, as well as a sixth subgroup of PD cases without variants in any PD-risk system (Other). The D, D/O and D/E subgroups represented the majority of individuals, as expected given the known involvement of DAn in PD. Beyond these cases, we noted a large minority of individuals assigned to the E and O-groups, comprising approximately 21.5% of PD (193/899). No differences in common risk were observed between subgroups, as quantified by polygenic risk scores of nominally significant common single nucleotide polymorphisms (p < 0.05; MAF > 0.05), although all subgroups exhibited higher PRS than controls (**Fig. 4b**).

**Figure 4.**
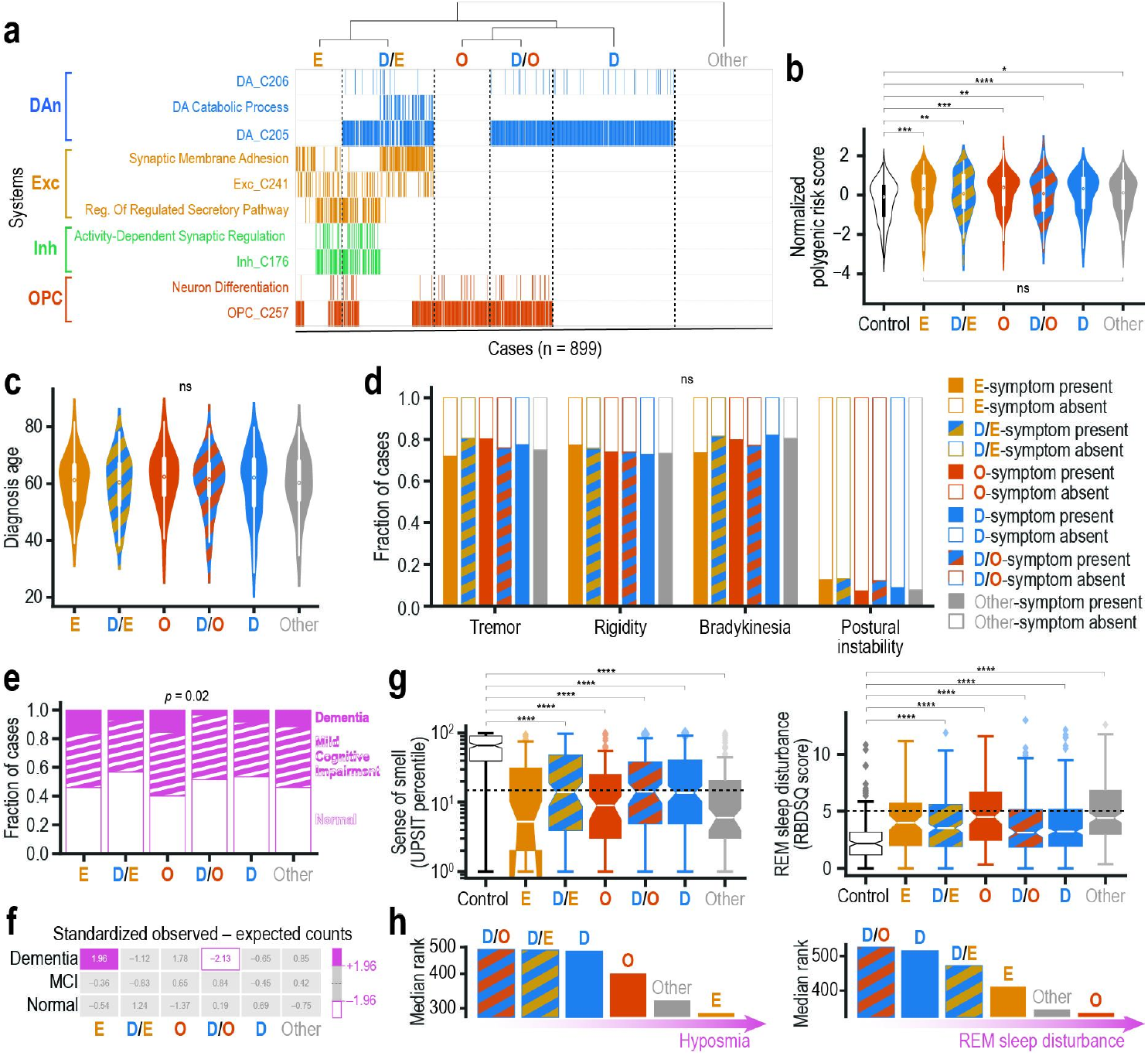
Classification of patient subgroups and associations with symptoms. (**a**) Clustering of PD cases (columns) into six subgroups (top dendrogram with labels) based on the presence or absence of minor allele variants in each of the ten PD-risk systems (rows). Systems are organized and colored by associated cell type: DAn (blue), Exc (gold), Inh (green), and OPC (orange). (**b**) Distribution of normalized polygenic risk scores (PRS) across groups. PRS were computed from GWAS summary statistics reported by Nalls et al. using SNPs with MAF > 0.05 and *p* < 0.05. Violin plots display score distributions; white dots denote medians and black bars interquartile ranges. *p < 0.05, **p < 0.01, ***p < 0.001, ****p < 0.0001; ns = not significant (two-tailed Mann-Whitney *U* test). (**c**) Ages at diagnosis for each PD subgroup. ns, not significant (two-tailed Mann-Whitney *U* test). (**d**) Motor symptoms at diagnosis for each PD subgroup. ns, not significant (Fisher’s exact test). (**e**) Distribution of cognitive states (normal; mild cognitive impairment or MCI; dementia) for each PD subgroup. *p* = 0.02 (chi-squared test). (**f**) Contingency table showing the standardized differences between observed and expected counts for each cognitive state (rows) across subgroups (columns). Purple indicates higher observed counts than expected, while white indicates lower observed counts, both outside the 95% confidence interval. (**g**) Left, sense of smell, measured as the University of Pennsylvania Smell Identification Test (UPSIT) percentile, across controls and the six subgroups, with the 15th percentile (dashed line) used as a cutoff for hyposmia. **** p < 0.0001 (two-tailed Mann-Whitney *U* test). Right, same for REM sleep behavior disturbance, assessed using the REM Sleep Behavior Disorder Screening Questionnaire (RBDSQ), with a score of five (dashed line) used as the threshold for probable RBD diagnosis. **** p < 0.0001 (two-tailed Mann-Whitney *U* test). (**h**) Median rank scores for hyposmia (left) and REM sleep disturbance (right) across subgroups. Lower ranks indicate stronger associations with the respective clinical features. PD, Parkinson’s Disease; DAn, dopaminergic neuron; Exc, excitatory neuron; Inh, inhibitory neuron; OPC, oligodendrocyte progenitor cell.

Next, we examined whether these six PD subgroups are predictive of clinical PD phenotypes. We found no differences across subgroups in age at diagnosis (**Fig. 4c**) or motor systems, such as tremor, rigidity, bradykinesia or postural instability (**Fig. 4d**). The subgroups did differ significantly in non-motor symptoms, however, with distinct distributions of normal cognition, mild cognitive impairment (MCI), and dementia (chi-squared test, *p* = 0.02; **Fig. 4e**). These observations could be attributed to an overrepresentation of cases with dementia in the E-group and an underrepresentation of cases with MCI in the D/O-group (**Fig. 4f**). For prodromal symptoms, all six subgroups scored significantly lower on the University of Pennsylvania Smell Identification Test (UPSIT) and higher on the Rapid Eye Movement (REM) Sleep Behavior Disorder Screening Questionnaire (RBDSQ) compared to non-PD controls (**Fig. 4g**; two-tailed Mann-Whitney *U* test, *p* < 0.0001), with the E-group showing the most severe hyposmia, and the O-group the most severe REM sleep disturbance (**Fig. 4h**; Kruskal-Wallis *H* test, *p* < 0.0001).

### An OPC system affecting myelin integrity

Of the PD risk systems used for disease classification, we further examined “OPC_C257,” which did not align with any known biological processes (**Fig. 3g,h**). This system was associated with oligodendrocyte precursor cells based on network proximity to documented OPC markers^15^ such as *TSC22D4* (Transforming growth factor β1 Stimulated Clone 22 Domain family 4), which plays a role in transcriptional regulation of myelin^18^.

Oligodendrocytes are uniquely responsible for myelin formation in the central nervous system. Therefore, we chose to interrogate this system using data from diffusion tensor imaging (DTI), a neuroimaging technique that measures the structural integrity of myelinated axons. DTI is based on the principle of fractional anisotropy (FA) in diffusion of water molecules: restricted diffusion along a preferred direction (high FA close to 1.0) indicates intact well-organized myelination, whereas diffusion less directional (low FA close to 0.0) indicates loss of myelin structural integrity. We examined DTI data of the substantia nigra from 152 post-mortem PD cases, of which 47 had genetic variants in the OPC_C257 system and 105 did not (**Fig. 5a**). We found that cases with OPC_C257 genetic variants displayed significant reductions in the measured FA, with differences observed on both the ipsilateral and contralateral sides of the brain and across the rostral, middle, and caudal regions of the substantia nigra (**Fig. 5b**).

**Figure 5.**
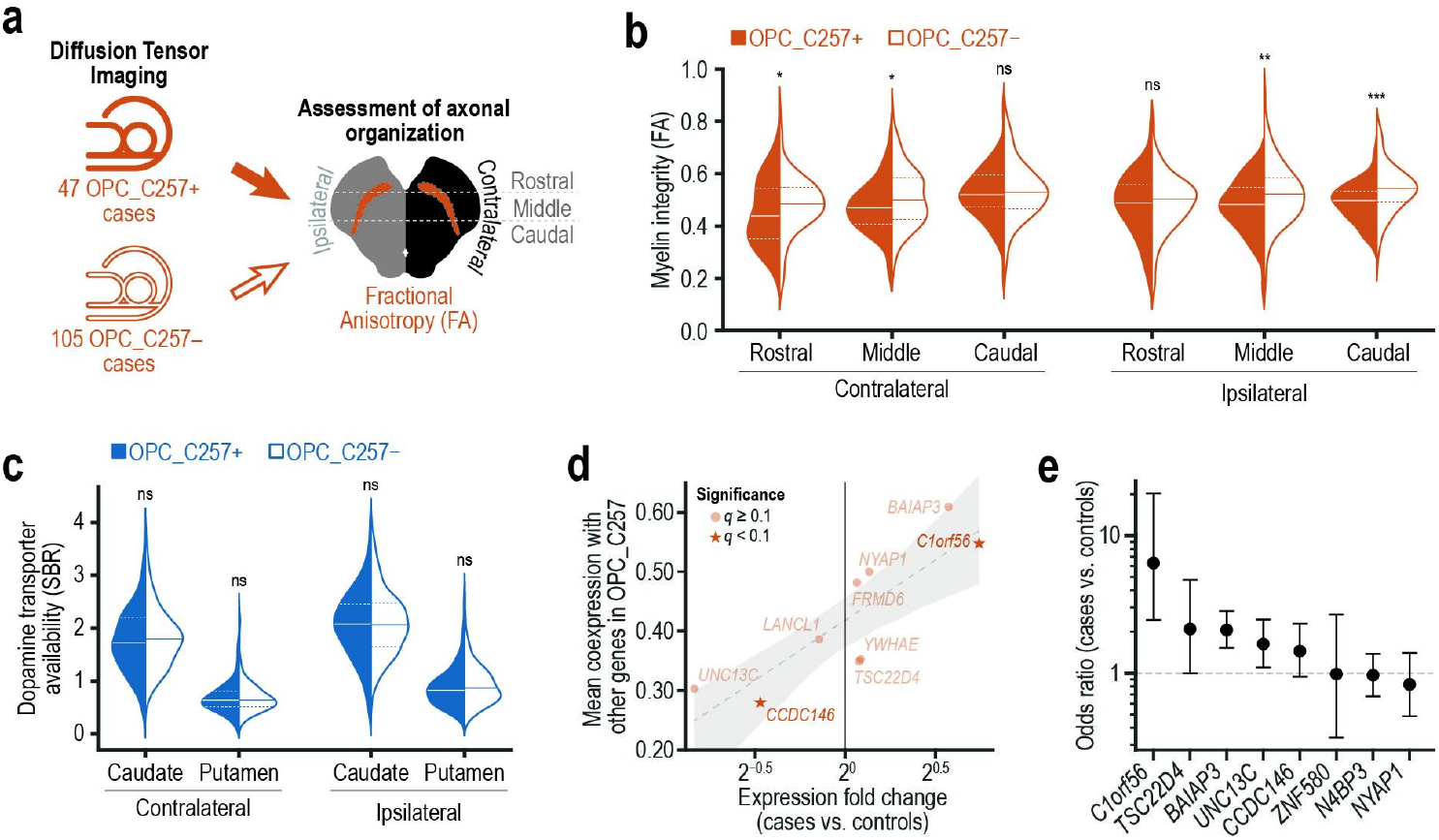
Analysis of a system (OPC_C257) linked to oligodendrocyte precursor cells. (**a**) Exploration of the OPC_C257 system using diffusion tensor imaging (DTI) data from 152 PPMI cases, comprising 47 with variants (OPC_C257+) and 105 without variants (OPC_C257−) in the system. Fractional anisotropy (FA) was measured on both the contralateral and ipsilateral sides of the rostral, middle, and caudal regions of the substantia nigra to evaluate white matter integrity. (**b**) Distribution of white matter integrity in OPC_C257+ (orange) and OPC_C257− cases (white) summarized by violin plots, with median (solid line) and interquartile range (dashed lines) indicated. FA values were normalized using the FA of a reference region for each individual. * *p* < 0.05, ** *p* < 0.01, *** *p* < 0.001; ns = not significant (one-tailed Mann-Whitney *U* test). (**c**) Distribution of dopamine transporter availability in OPC_C257+ (blue) OPC_C257− summarized as violin plots, with median (solid line) and interquartile range (dashed lines) indicated. Striatal binding ratio (SBR) values were calculated for the four striatal regions by extracting count densities and normalizing them to the count density of the reference region. ns = not significant (two-tailed Mann-Whitney *U* test). (**d**) Scatterplot of OPC_C257 genes, plotted by their mean coexpression in OPC cells when paired against each other gene in the system (y-axis; absolute Spearman correlation) versus their fold change in expression between cases and controls (x-axis). Genes with *q* < 0.1 from differential expression analysis are marked with a star (☆). A regression line (dashed line; slope = 0.2, R^2^ = 0.71, *p* = 0.004) is shown with a 95% confidence interval (gray shading). (**e**) Forest plot showing the strength of PD association for each gene in the OPC_C257 system. Odds ratios >1 indicate a stronger association with cases, whereas odds ratios <1 indicate a stronger association with controls. Error bars represent 95% confidence intervals.

To test whether the observed myelin deficits were secondary to dopaminergic degeneration, we examined dopamine transporter (DAT) scans acquired using single-photon emission computed tomography. These scans use [^123^I]ioflupane, a radiotracer that binds specifically to DAT, to measure its availability in the striatum as a proxy for DAn integrity. We found similar DAT levels in the caudate and putamen between variant carriers and non-carriers bilaterally (**Fig. 5c**), supporting a distinct, non-dopaminergic mechanism of pathology. The expression changes observed in PD align with native coexpression patterns of these genes in OPCs (**Fig. 5d**), suggesting that they constitute a coordinated gene network that becomes dysregulated in PD, rather than a random set of unrelated genes. Four genes showed moderate to high associations with PD risk (**Fig. 5e)**: *C1orf56* (OR 6.3), *TSC22D4* (OR 2.1), BAIAP3 (OR 2.1), and UNC13C (OR 1.6).

### Structural and functional characterization of C1orf56 variants

Notably, *C1orf56* (chromosome 1 open reading frame 56) is a largely uncharacterized gene with no prior link to PD or known role in the brain. It harbored four missense variants specific to OPC_C257+ cases, all located within an epigenetically modified accessible region (**Fig. 6a**).

**Figure 6.**
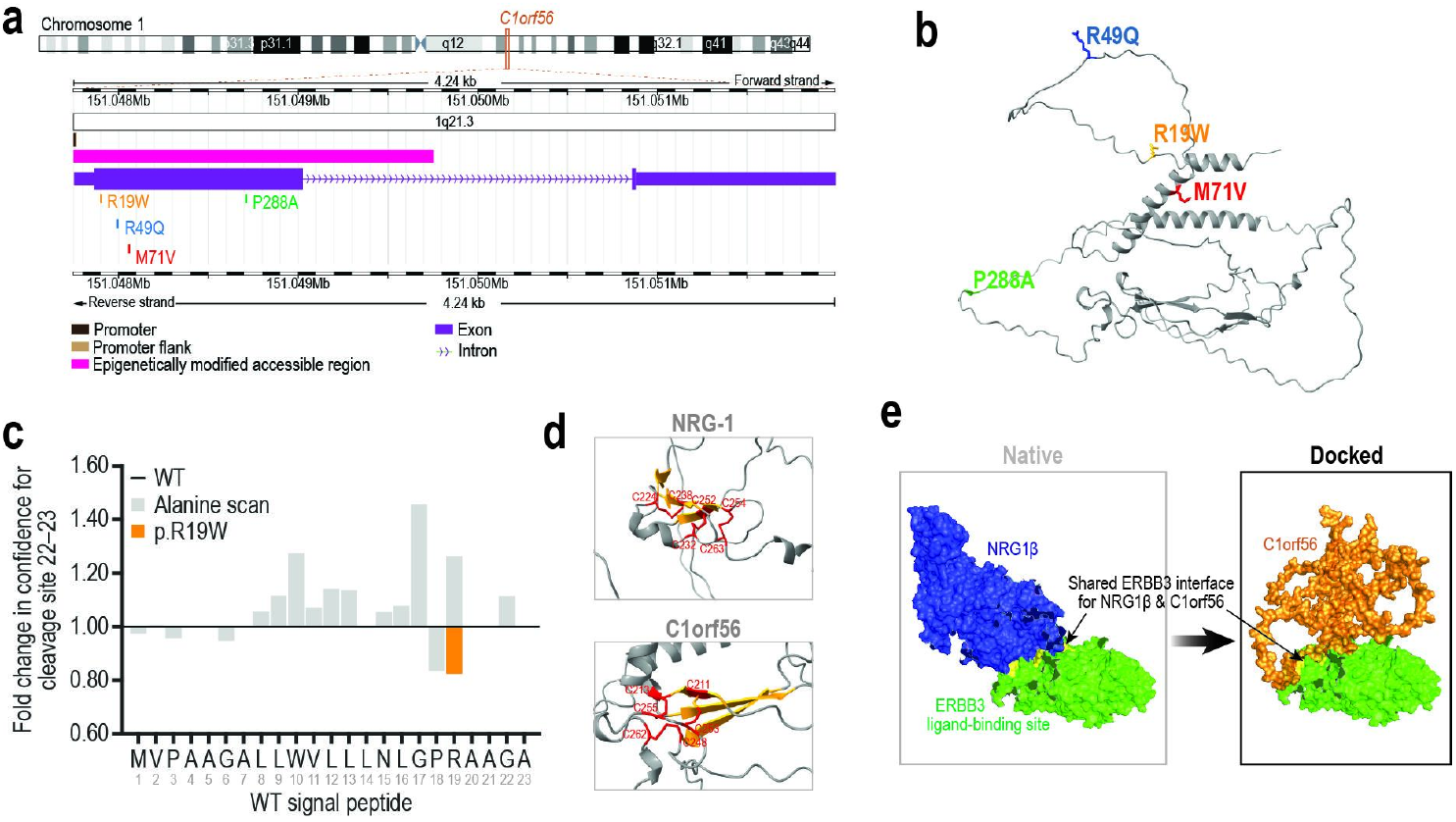
Structural and functional characterization of C1orf56 and associated variants. (**a**) Genomic view of *C1orf56*, showing the location of four missense variants identified in OPC_C257+ cases. A schematic of chromosome 1 highlights the cytogenetic band (orange) 1q21.3 that contains *C1orf56* gene. The zoomed-in window spans approximately 4.24kb. Exons (purple rectangles) are connected by an intron (arrowed line, arrows denote transcriptional orientation of the forward strand). Regulatory annotations are shown below: promoter (black), promoter flank (brown), and epigenetically accessible region (magenta). Four missense variants identified in OPC_C257+ cases are marked by their dbSNP identifiers (rs74856367, rs75577601, rs1132889, rs143972785). (**b**) AlphaFold predicted structure of the C1orf56 protein (gray) with four missense variants highlighted (red, orange, green, and blue). (**c**) Fold change in confidence for the predicted signal peptidase cleavage site at positions 22-23 relative to wild type (WT, black line) as determined by Signal P 6.0. Gray bars show results from alanine substitutions at each residue across the signal peptide sequence (MVPAA…AGA), while the orange bar indicates the naturally occurring variant rs7485637. Values above 1.0 represent increased cleavage site confidence, while values below 1.0 represent reduced confidence compared to WT. (**d**) Enlarged view of the EGF-like fold for Neuregulin-1 (NRG-1; top) and C1orf56 (bottom), showing a compact structure with an antiparallel β-sheet (yellow) stabilized by three disulfide bridges (∼2.0 Å) formed by six cysteines (red). (**e**) Predicted binding of C1orf56 to the ERBB3 receptor at the neuregulin-1 (NRG1β) interaction site. Left, Native complex showing NRG1β bound to the extracellular domain of ERBB3 (green). Right, Docked model of C1orf56 (orange) bound to ERBB3, highlighting the shared ERBB3 interface with NRG1β. Surface representations generated in PyMOL based on AlphaFold prediction for C1orf56 and docking models computed by HADDOCK.

The rs143972785 (p.M71V) variant lies within an α-helix (**Fig. 6b**) and was predicted to be structurally neutral (ΔΔG = –0.06 kcal/mol), while the oxidized-Met proxy (p.M71Q) produced a small stabilizing shift (ΔΔG = +0.79 kcal/mol). Although the variant is predicted to be benign (CADD PHRED = 2.9), its location within an α-helix and structural modeling suggest it could still affect function through loss of redox-sensitive behavior. The rs74856367 (p.R19W) variant within the signal peptide reduced the predicted confidence of cleavage at site 22-23 compared to wild type and alanine scans (**Fig. 6c**). This result aligns with its predicted deleteriousness (CADD PHRED = 16.7) and suggests impaired secretion and signaling. To further investigate the function of C1orf56, we next examined its predicted structure using AlphaFold^19^. We found that C1orf56 shares a predicted epidermal growth factor (EGF)-like fold with neuregulin-1 (**Fig. 6d**), characterized by an antiparallel β-sheet stabilized by three disulfide bridges formed by six cysteines. Neuregulins are secreted EGF-family ligands that bind to EGFRs, which regulate proliferation, differentiation, and, within the nervous system, glial development and myelination. Notably, C1orf56 is also predicted to be secreted and to bind at the same site as the neuregulin-1 beta isoform (NRG1β) on an EGFR family member, such as epidermal growth factor receptor 3 (ERBB3) with favorable energetics (**Fig. 6e**; HADDOCK score^20^ = −103, buried surface area = ∼1970 Å^2^), supporting a physical interaction. Together, these structural and functional predictions suggest that C1orf56 may regulate myelination through mechanisms similar to those of NRG1.

## DISCUSSION

Here we have shown that the genetic variants carrying risk for PD converge on genetic networks associated with four distinct cell types (**Fig. 3g**). Apart from DAn, our results implicate excitatory neurons, inhibitory neurons, and OPCs as primary and independent contributors to disease. In our study, astrocytes and microglia markers did not colocalize with common PD risk genes in the genetic network (**Fig. 2e**). These genetic findings contrast somewhat with results from single cell transcriptomics, which have reported gene expression changes associated with microglia and astrocytes in postmortem PD brains^12,13^. The lack of colocalization suggests that they may instead contribute through non-genetic mechanisms, given their well-established roles in neuroinflammation^21^, rather than serving as primary mediators of common genetic risk^11^.

Using the cell-type systems framework (**Fig. 3g**), we classified PD cases into six subgroups, each characterized by distinct genetic networks linked to specific PD symptoms (**Fig. 4a**). Given that not all subgroups harbor variants in DA systems, it is noteworthy that the non-canonical cases (E-group, OPC-group, Other) still develop motor symptoms such as tremor, rigidity, bradykinesia, and postural instability (**Fig. 4c**). Such individuals also experience significant non-motor symptoms, such as cognitive decline (**Fig. 4e**), hyposmia and REM sleep behavior disturbance (**Fig. 4g**). In contrast, individuals with DA system variants (D-, D/E- and D/O-group) exhibit strong motor symptoms (**Fig. 4c**) but mild non-motor symptoms (**Fig. 4g**), despite some of these individuals also carrying variants in non-DA systems (D/E- and D/O-group). In these mixed cases, it is possible that compromised DAn, primarily located in the substantia nigra and ventral tegmental area, compensate for the non-motor symptoms through their projections to brain regions, such as the prefrontal cortex^22–24^, hippocampus^25,26^, olfactory bulb^27^, or ventral periaqueductal gray^28^, which, in fact, regulate cognition, sleep and olfaction.

We found that individuals of the E-group, with variants in excitatory but not DAn systems, were significantly enriched for dementia (**Fig. 4d, e**). One excitatory neuron system, “Exc_C241” (**Fig. 3g**), was enriched with genes known to cause epilepsy^29^ (*q* = 3.4×10^−8^), including *CDKL5*^*30*,*31*^, *PCDH19*^*32*,*33*^, *SCN1A*^*34*,*35*^, and *LGI1*^*36*,*37*^. This enrichment points to a shared pathogenic pathway between PD and epilepsy, further supported by the incidence of motor symptoms in both diseases including dystonia, myoclonus, ataxia, tremor, and hypokinesia^38^. Furthermore, PD cases are nearly ten times more likely than controls to have had epilepsy in the two years before diagnosis^39^. Taken together, these genetic and clinical links suggest that disrupted excitation/inhibition balance may precede motor symptoms, and that antiepileptic drugs could potentially be repurposed for PD prevention, targeting E-group individuals^40^.

We identified an OPC-associated risk system of 11 genes (OPC_C257, **Fig. 3g**). Variants in this system were associated with reduced myelin integrity (**Fig. 5b**), in line with impaired maturation observed in patient-derived pre-oligodendrocytes^41^. Indeed, oligodendrocytes are only recently emerging as key contributors to PD pathology^42,43^. A marked reduction in oligodendrocytes has also been reported in the midbrain^12^, particularly within the substantia nigra pars compacta^13,44^. Whereas prior studies identified oligodendrocyte-specific expression patterns enriched for common genetic risk^11,45^, our findings integrate genetic, cellular, and phenotypic evidence to implicate OPC-related genes in linking myelin loss to non-motor symptoms.

*C1orf56*, a poorly characterized gene, emerges as the highest-risk gene within this system. We discovered that its gene product shares structural and functional features with NRG1, including a signal peptide for secretion (**Fig. 6c**), an EGF-like fold (**Fig. 6d**), and predicted EGFR binding (**Fig. 6e**). Notably, NRG1 signaling can delay or fine-tune OPC differentiation and promote proliferation and survival during development^46,47^ or in response to injury. Variants in NRG1, such as rs2439272, rs6988339^48^, rs6994992^49^, have been associated with white matter changes^49^, as well as with neurodevelopmental psychiatric disorders, such as schizophrenia^50^ and attention deficit hyperactivity disorder^51^. Taken together, these findings support the role of C1orf56 as a signaling protein, possibly involved in OPC differentiation and myelination. Accordingly, we hereafter refer to this gene as Neuregulin-6 (*NRG6*), the sixth protein identified with neuregulin-like properties^52^.

*NRG6* was found to be associated with four missense variants (**Fig. 6a**). We delineated potential mechanisms for two of these variants, with one located in the α-helix (rs143972785; **Fig. 6b**) and the other in the signal peptide (rs74856367; **Fig. 6c**). While p.M71V (rs143972785) is not a grossly destabilizing switch, it removes an oxidation-sensitive regulatory site, potentially impairing the ability of NRG6 to respond to oxidative stress, a well-established contributor to PD ^53,54^. p.R19W (rs74856367) significantly lowers the confidence of cleavage site recognition compared to wildtype and alanine controls (**Fig. 6c**), suggesting impaired secretion of NRG6 into the extracellular space.

In summary, our work has highlighted the role of cell types beyond DAn in PD, particularly OPCs and their impact on myelin. Although this multi-cellular dimension of PD introduces additional complexity, it also opens avenues for novel therapeutic targets and the potential repurposing of existing neuropsychiatric drugs for prevention. We expect that the network-based patient stratification implemented here can be readily generalized to other neurodegenerative diseases traditionally examined through selective vulnerability, such as Alzheimer’s disease^55^, multiple sclerosis^56^, amyotrophic lateral sclerosis, and Huntington’s disease, creating the potential for individualized treatments based on dysregulated cell types.

## METHODS

### Data acquisition

The HumanNet-XC v3 network^16^ was retrieved from the ndexbio.org (uuid: 8f929fb5-3ac6-11ed-b7d0-0ac135e8bacf). This network combines inferred co-functional relationships from multiple datasets, including PubMed co-citation, gene co-expression, protein-protein interactions, genetic interactions, protein domain co-occurrence, and genomic context similarity. Network nodes were filtered to include only genes expressed in the human brain proteome (n = 15,331), as defined in the Human Protein Atlas^57,58^. The largest connected component of the resulting subnetwork was used for further analysis (18,593 genes, 1,125,494 gene-gene interactions). 1,125,494 functional relationships among 18,593 human genes.

We obtained GWAS summary statistics from Nalls et al.^14^, the largest meta-analysis of PD based on 37,688 cases, 18,618 proxy cases (having a first degree relative with PD), and approximately 1.4 million controls. A total of 90 genome-wide significant genetic variants (p < 5×10^−8^) spanning 78 genomic regions were mapped to their nearest genes, resulting in 86 distinct genes associated with common risk for PD.

For cell markers, data for the eight cell types were downloaded from Cell Marker 2.0, a manually curated database that integrates data from single-cell sequencing and other human and mouse resources (see **Table S2** for search keywords). Only distinctive markers were retained for each cell type (see **Table S3** for finalized gene sets used for network propagation). HumanNet contained most PD risk genes (73/86) and distinct cell markers: 16/18 DAn, 27/28 excitatory neuron, 28/28 inhibitory neuron, 94/96 OPC, 293/328 microglia, 380/394 astrocyte, 19/32 hepatocyte, and 11/15 podocyte markers.

### Network propagation and colocalization

Common risk genes and cell markers were propagated and colocalized in HumanNet-XC^16^ using the Python package NetColoc 1.0 (https://github.com/ucsd-ccbb/NetColoc/)^59,60^. Briefly, a Random Walk with Restart algorithm^61^ was used to model propagation of heat (also called heat diffusion) along the edges in the network, where heat diffuses from an initial seed of “hot” genes to its neighbors. A constant fraction of heat dissipates from each gene per iteration, stabilizing at the closed-form solution in Equation 1:

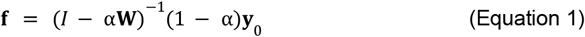

where **f** is the stable vector of heat values for each node, **y**_0_is the vector of seed genes, **W** is the column-normalized adjacency matrix of the network (here, HumanNet), and α ∈ (0, 1) is the dissipation constant which was set to α = 0.5. Following propagation, a network proximity score (NPS) was calculated for each gene g by comparing the observed heat values to the null distribution created from 1,000 random sets of initial seed genes, which were generated to maintain the same size and degree distribution as the original seed set as described by Guney et al.^62^ For each seed set *S*, the NPS was expressed as a z-score by comparing *F* _g, *s*_, the observed heat at g, versus 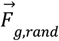, the mean heat from the null distribution, as described in Equation 2:

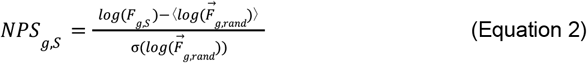

where ⟨⟩ represents the mean of a vector, and σ represents its standard deviation. All heat values were log-transformed (ln) to approximate a normal distribution.

NPS was calculated separately for the common risk genes (NPS_PD_) and for the sets of cell markers across each of eight cell types (NPS_cell_type_). To identify genes proximal to both gene sets, the combined NPS (NPS_both_) was calculated as the product:

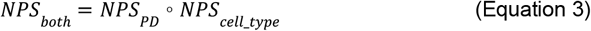

These scores were then thresholded to define a colocalized gene set: NPS_cell_type_ > 1.5, NPS_PD_ > 1.5, NPS_both_ > 1.5.

### Identification of molecular systems in distinct cell types

Each colocalized gene set (see above) was analyzed to discover its network community substructure using the Hierarchical community Decoding Framework (HiDeF)^63^ implemented in the Community Detection Application and Service (CDAPS) Python package^64^. HiDeF leverages persistent homology to identify densely interacting network community structures across scales. Briefly, HiDeF clusters a network at various modularity resolutions using the Louvain algorithm, where higher resolutions produce smaller communities, and lower resolutions produce larger ones. Persistent communities are identified as those repeatedly detected across resolutions, then organized hierarchically. We ran HiDeF with a maximum resolution of 5 and default settings for other parameters. Resulting communities (systems) were labeled with informative biological names using Enrichr^65^ to identify enriched GO Biological Processes (FDR-corrected using the Benjamini-Hochberg procedure, *p* < 0.05), followed by adjustment of system labels through expert curation. For further analysis we focused on only those systems at the leaves of the HiDeF hierarchy to arrive at a distinct set of compact systems. All subsequent network figures were created using Cytoscape. Spring-embedded layout was used for subnetwork figures and manually adjusted (**Fig. 1a,3a,3g**).

### Variant analysis of network systems in independent samples

Whole-genome sequencing data for ‘Healthy Control,’ ‘Parkinson’s Disease,’ and ‘Prodromal’ cohorts was downloaded from PPMI^66^ (http://www.ppmi-info.org) as VCF (Variant Call Format) files. Briefly, previous work by PPMI had sequenced whole-blood DNA samples on the Illumina HiSeq X platform and aligned these sequences to the GRCh38DH reference genome. Variants were called using GATK Best Practices^67^ and Variant Quality Score Recalibration^68^. To annotate and standardize the variant data for downstream analysis, we converted VCF to MAF (Mutation Annotation Format) files^69^, ensuring consistent annotation with Ensembl’s Variant Effect Predictor (VEP)^70^. Coding variants were filtered by selecting those classified as “Frame_Shift_Del,” “Frame_Shift_Ins,” “Splice_Region,” “Splice_Site,” “Missense_Mutation,” “Nonsense_Mutation,” and “Nonstop_Mutation.”

For each system *s*, the system mutation burden *x*_*i*_ in individual *i* was computed as the aggregate number of variants detected within the genes of that system for that individual. The corresponding case/control status (*y*_*i*_) was modeled as a function of the burden score using logistic regression:

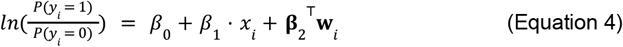

where *P* (*y*_*i*_ = 1) is the probability that individual i is a case, β_0_is the intercept, β_1_ is the effect size of the burden score, **W**_*i*_ represents the vector of covariates, including age, sex and the first two principal components to account for confounding factors, and **β**_2_ is the vector of regression coefficient for the covariates. After fitting of parameters, the significance of the burden effect (*β*_*1*_ ≠ 0) was evaluated using the Wald test. The odds ratio (OR) for the burden score is given by:

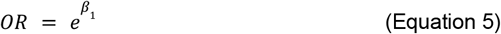

which quantifies how each additional variant in the system alters the odds being a case.

### Single-cell transcriptomic analysis

Single-cell RNA-seq data from post-mortem midbrain samples of six healthy controls and five idiopathic Parkinson’s disease patients were obtained from Smajicć et al^12^. Unique molecular identifier (UMI) counts were normalized to adjust for sequencing depth, ensuring gene expression levels were comparable across cells, and then log_2_ normalized. Genes with expression ≥ 0.1 were retained. To assess co-expression between gene pairs, cosine similarity was computed between their expression vectors across all cells of the same cell type. Relevant to **Figs. 3E, 3F, 5A, 5E**.

### Patient stratification by cell type and symptom profiling

The ten systems with an odds ratio > 1.5 (see above) were grouped by cell type of origin: DAn, excitatory neuron, inhibitory neuron, or OPC. Hierarchical clustering of variant patterns across the matrix of systems versus PD cases (rows vs. columns) was performed using Ward’s method^71^, with six clusters of PD cases formed by cutting the dendrogram at a distance threshold of 10. Cluster order was optimized through optimal leaf ordering^72^ for better visualization. To assess clustering robustness, we performed bootstrap resampling^73^: In each of 1,000 bootstrap iterations, a new dataset was generated by sampling individuals with replacement, hierarchical clustering was re-applied using a fixed distance threshold of 10, and the resulting cluster assignments were compared to the original clusters. Cluster stability was quantified using the Jaccard Similarity Index (JSI), recording the highest JSI for each original cluster in each iteration.

To identify differences in phenotypes, we downloaded age and motor symptoms– tremor, rigidity, bradykinesia, and postural instability– at diagnosis from PPMI. For non-motor symptoms, we considered cognitive decline and two prodromal symptoms, hyposmia and REM sleep behavior disturbance (RBD). Cognitive state (COGSTATE) values were obtained for each individual, classified as normal (1), mild cognitive impairment (2), and dementia (3). The highest COGSTATE value recorded for an individual across all visits was used to determine if an individual had developed cognitive decline and, if so, the severity. Hyposmia was assessed using the age- and sex-adjusted UPSIT percentile values, with the 15th percentile used as a cutoff, consistent with prior studies^74,75^. RBD was assessed as the average RBDSQ score across visits, as RBD symptoms do not typically worsen over time^76^. An RBDSQ score ≤ 5 was used as the threshold for probable RBD diagnosis, consistent with prior studies^77–79^.

### Diffusion Tensor Imaging (DTI) analysis

We obtained fractional anisotropy (FA) values for the rostral, middle, and caudal regions of the substantia nigra from PPMI^66^, processed per Schuff et al^80^. Briefly, DTI data were acquired using a cardiac-gated two-dimensional single-shot echo-planar sequence to map brain water diffusion, with standard scalar measures, including FA, and vector-valued main directional diffusivity derived from the diffusion tensor^80^. Cases with unilateral symptoms (n = 153) were analyzed to map left and right sides as ipsilateral or contralateral. FA values for each side were normalized to the corresponding reference regions in the cerebral peduncle region^80^ for each individual. Cases were classified into OPC_C257+ and OPC_C257– subtypes based on the presence or absence of variants in any of the 11 genes in the OPC_C257 system, and their normalized FA values were compared using a one-tailed Mann-Whitney *U* test.

### C1orf56 structural and docking analysis

The three-dimensional structure of the C1orf56 protein was obtained from the AlphaFold Protein Structure Database (https://alphafold.ebi.ac.uk). The predicted model was downloaded in PDB format and manually inspected using PyMOL (v2.5.4, Schrödinger, LLC) to assess EGF-like domain architecture, including the presence of β-sheets and disulfide bridges between conserved cysteine residues.

To assess potential binding between C1orf56 and the ERBB3 receptor, protein–protein docking simulations were performed using the HADDOCK 2.4 web server (https://wenmr.science.uu.nl/haddock2.4/)^81^. The crystal structure of the extracellular domain of ERBB3 in complex with NRG1β was retrieved from the Protein Data Bank (PDB ID: 3U7U). Chains corresponding to ERBB3 and NRG1β were retained, and all other chains and ligands were removed. The AlphaFold-predicted structure of C1orf56 was used as the second binding partner. Docking was performed using default HADDOCK parameters. Active residues for NRG1β were defined based on its known interaction interface with ERBB3 and included residues 252, 254, 257, 264, 265, 266, 267, 270, 293, 294, 295, and 457. For C1orf56, active residues were selected based on surface accessibility within its predicted EGF-like domain and included residues 248–274. The resulting docking models were ranked by HADDOCK score, and the top-scoring cluster was selected for further analysis.

All structural figures and molecular surface representations (**Fig. 5g,h**) were generated using PyMOL. Docked complexes were visualized to compare the binding orientation of C1orf56 with that of NRG1β and to illustrate potential receptor engagement through the shared extracellular domain of ERBB3.

## RESOURCE AVAILABILITY

All colocalized networks and hierarchies generated in this study are available on NDEx as part of the network set. DOIs will be issued upon publication.

## ACKNOWLEDGEMENTS

This work was supported by the following grants: NIDA P50 DA037844 from the National Institutes of Health to T.I.

Data used in the preparation of this article was obtained on [2024-07-29] from the Parkinson’s Progression Markers Initiative (PPMI) database (www.ppmi-info.org/access-dataspecimens/download-data), RRID:SCR_006431. PPMI is a public-private partnership funded by the Michael J. Fox Foundation for Parkinson’s Research and funding partners, including 4D Pharma, Abbvie, AcureX, Allergan, Amathus Therapeutics, Aligning Science Across Parkinson’s, AskBio, Avid Radiopharmaceuticals, BIAL, BioArctic, Biogen, Biohaven, BioLegend, BlueRock Therapeutics, Bristol-Myers Squibb, Calico Labs, Capsida Biotherapeutics, Celgene, Cerevel Therapeutics, Coave Therapeutics, DaCapo Brainscience, Denali, Edmond J. Safra Foundation, Eli Lilly, Gain Therapeutics, GE HealthCare, Genentech, GSK, Golub Capital, Handl Therapeutics, Insitro, Jazz Pharmaceuticals, Johnson & Johnson Innovative Medicine, Lundbeck, Merck, Meso Scale Discovery, Mission Therapeutics, Neurocrine Biosciences, Neuron23, Neuropore, Pfizer, Piramal, Prevail Therapeutics, Roche, Sanofi, Servier, Sun Pharma Advanced Research Company, Takeda, Teva, UCB, Vanqua Bio, Verily, Voyager Therapeutics, the Weston Family Foundation and Yumanity Therapeutics.

The FP7 WeNMR (project# 261572), H2020 West-Life (project# 675858), the EOSC-hub (project# 777536) and the EGI-ACE (project# 101017567) European e-Infrastructure projects are acknowledged for the use of their web portals, which make use of the EGI infrastructure with the dedicated support of CESNET-MCC, INFN-LNL-2, NCG-INGRID-PT, TW-NCHC, CESGA, IFCA-LCG2, UA-BITP, TR-FC1-ULAKBIM, CSTCLOUD-EGI, IN2P3-CPPM, SURFsara and NIKHEF (NWO projects# 10236 and 17437), and the additional support of the national GRID Initiatives of Belgium, France, Italy, Germany, the Netherlands, Poland, Portugal, Spain, UK, Taiwan and the US Open Science Grid.

Finally, we thank Steven A. Prescott and Tomasz J. Nowakowski for reviewing the manuscript.

## SUPPLEMENTAL INFORMATION

Figure S1 and Tables S1-3

