## Supplemental information for "Redefining Parkinson’s Disease by Dysregulated Genetic Networks in Distinct Cell Types"

**Table S1.** The 22 systems significantly associated with case/control status in the PPMI cohort based on system-wide burden modeling

| ID | Cell_type | Name | Genes | OR | CI_OR_low | CI_OR_high | p_val | FDR adjusted_p |
|---|---|---|---|---|---|---|---|---|
| DA_C212 | DA | DA_C212 | UTP25,WLS,DCAF16,EN1 | 0.366623797 | 0.252759354 | 0.523870029 | 6.35E-08 | 1.76E-06 |
| DA_C206 | DA | DA_C206 | CACNB3,UBIAD1,GNG8,SYT1,GNB4 | 4.30332214 | 2.178899813 | 9.286771641 | 6.81E-05 | 0.000686955 |
| DA_C205 | DA | DA_C205 | RAB32,C12orf29,REM2,LRRK2,RIT2 | 2.075896789 | 1.765834172 | 2.447903953 | 1.79E-18 | 1.99E-16 |
| DA_C195 | DA | DA_C195 | LAMB2,CPE,PTPRO,INF2,WDR73,CPO,CD2AP,LMX1B,SCARB2 | 1.231634074 | 1.106468505 | 1.372666595 | 0.000150141 | 0.001190407 |
| DA_C203 | DA | DA_C203 | EPB41L2,SHANK2,ZBTB20,AUTS2,EPB41L3,ZSCAN1 | 0.781766414 | 0.680544412 | 0.896837674 | 0.000465916 | 0.003042157 |
| DA_C209 | DA | DA Catabolic Process | MAOB,GRIN2B,NETO1,MAOA | 3.56323 0247 | 2.009474375 | 6.765302972 | 3.64E-05 | 0.000403502 |
| DA_C208 | DA | DA_C208 | CRHR1,PKNOX2,ZFHX4,MAB21L1,TTC12 | 0.788472684 | 0.669463865 | 0.927203543 | 0.004203621 | 0.02028704 |
| Exc_C241 | Exc | Exc_C241 | CDKL5,PCDH19,SCN1A,WSCD1,LGI1 | 2.386916325 | 1.642141329 | 3.565517179 | 1.02E-05 | 0.00016154 |
| Exc_C243 | Exc | Exc_C243 | PTPRF,GRIA1,NETO1,GRIN2B,LRFN4 | 2.770258017 | 1.948189205 | 4.033208585 | 3.75E-08 | 1.39E-06 |
| Exc_C235 | Exc | Exc_C235 | TOX,PRKAR2B,TCF7,IKZF2,IKZF1,SCNM1,TOX2 | 1.27677932 | 1.091620879 | 1.496079186 | 0.002356938 | 0.013081007 |
| Exc_C233 | Exc | Regulation Of Regulated Secretory Pathway | SYN1,SYT6,SYT1,SYT5,CHGB,SYP,SYT4,VSNL1 | 1.695166757 | 1.306380849 | 2.244473617 | 0.000124371 | 0.001078233 |
| Exc_C242 | Exc | Exc_C242 | QPRT,CREB1,DEAF1,ADNP,TLL1 | 0.519393931 | 0.344743555 | 0.768022186 | 0.00129578 | 0.007570086 |
| Inh_C177 | Inh | Inh_C177 | KLF3,LHX6,ONECUT2,ZNF76,MAP2,CUX2 | 0.69900501 | 0.591278553 | 0.824348735 | 2.36E-05 | 0.000291601 |
| Inh_C176 | Inh | Inh_C176 | CHGB,SYT4,SYN1,SYP,RPRD1B,SYT6 | 1.656062789 | 1.27334894 | 2.196396509 | 0.000271819 | 0.001885746 |
| Inh_C178 | Inh | Inh_C178 | PSEN1,CA3,CPE,TG,GCA,CGA | 1.495926593 | 1.275074476 | 1.759677087 | 9.38E-07 | 1.74E-05 |
| Inh_C182 | Inh | Inh_C182 | DPP10,DLG2,SHANK2,PLXNA4,DYNLL2 | 0.775436881 | 0.680475434 | 0.882727822 | 0.000126279 | 0.001078233 |
| Inh_C187 | Inh | Activity-De | SCN1A,RASD1,REM2,PCDH19 | 4.8181 | 2.44795 | 10.5943 | 2.09E | 0.0002896 |

| | | | | | | | | |
|---|---|---|---|---|---|---|---|---|
| | | pendent Synaptic Regulation | | 346 | 5103 | 5522 | -05 | 92 |
| OPC_C283 | OPC | OPC_C283 | LHX6,NHLH2,ZNF76,STMN3 | 0.624479798 | 0.437790831 | 0.881281121 | 0.008150231 | 0.036187024 |
| OPC_C256 | OPC | OPC_C256 | SHANK2,NRXN2,EPB41L1,EPB41L2,LRRTM2,PDZD2,GPATCH8,DLG2,DLGAP4,EPB41L3,DLGAP3 | 0.875336507 | 0.802523143 | 0.954203369 | 0.00255622 | 0.013511449 |
| OPC_C257 | OPC | OPC_C257 | YWHAE,TSC22D4,FRMD6,BAIAP3,CCDC146,LANCL1,N4BP3,C1orf56,ZNF580,NYAP1,UNC13C | 1.540790931 | 1.315839945 | 1.811529087 | 1.13E-07 | 2.51E-06 |
| OPC_C265 | OPC | OPC_C265 | KAZALD1,MDGA1,PPIP5K2,C1orf162,NRN1,NTNG1,THY1,BST1,IP6K1 | 0.762356134 | 0.648777704 | 0.893930398 | 0.000897257 | 0.005533086 |
| OPC_C278 | OPC | Neuron Differentiation | EN1,LMX1B,CNPY1,FGF20,PITX3 | 2.380656346 | 1.367373408 | 4.303779591 | 0.002855229 | 0.014405927 |

DA, dopaminergic neurons; Exc, excitatory neurons; Inh, inhibitory neurons; OPC, oligodendrocyte progenitor cells

**Table S2.** Keywords used for each cell type in Cell Marker 2.0

| Cell type | Keywords |
|---|---|
| Dopaminergic neuron | “Dopaminergic neuron,” “Midbrain dopaminergic neuron,” and “Dopamine neuron” |
| Excitatory neuron | ‘Glutamatergic neuron,” “Excitatory neuron,” “Broad excitatory neuron,” “Superficial layer excitatory neuron,” and “Deep layer excitatory neuron” |
| Inhibitory neuron | “GABAergic neuron” and “Inhibitory neuron” |
| Astrocyte | “Astrocyte” |
| Microglia | “Microglial cell,” “Microglial-specific cell,” and “Microglia-like cell” |
| Oligodendrocyte progenitor cell | “Oligodendrocyte progenitor cell” and “Oligodendrocyte precursor cell” |
| Hepatocyte | “Hepatocyte” |
| Podocyte | “Podocyte” |

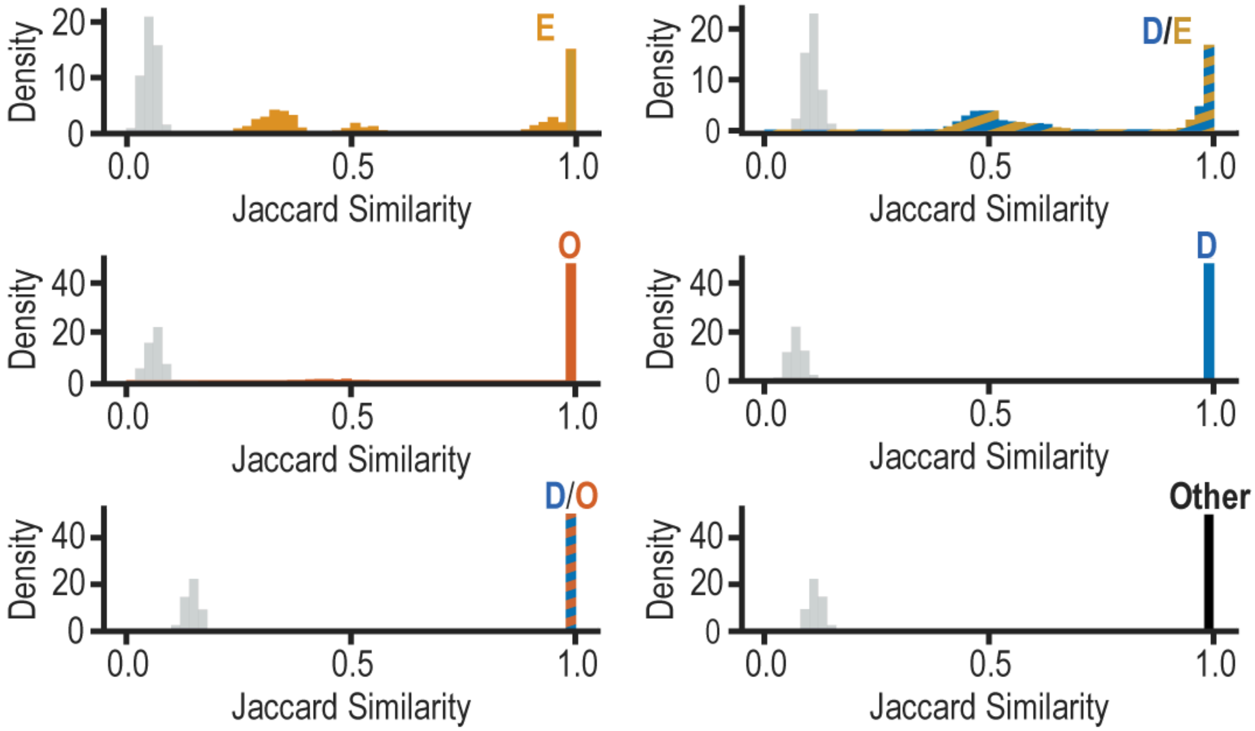


**Figure S1. Cluster robustness analysis of patient subgroups.** Density plots showing the distribution of observed Jaccard Similarity Index (JSI) values across bootstrap samples for each patient (colored distributions) alongside the null distribution of JSI values (gray), which serves as a baseline for comparison (see **Methods**).
